# Eomesodermin defines uterine NK cells crucial for pregnancy success in mice

**DOI:** 10.1101/2025.03.29.645626

**Authors:** Josselyn D. Barahona, Liping Yang, Wayne M. Yokoyama

## Abstract

Uterine natural killer cells have been thought to be critical for reproductive success, and their developmental origins remain unclear. Here, we demonstrate that Eomesodermin is a key transcription factor determining the lineage of tissue-resident NK cells within the uterus both at steady-state and during pregnancy. Ablation of Eomesodermin in Ncr1-expressing cells results in the loss of tissue-resident NK cells in both the virgin and pregnant uterus, suggesting that uterine tissue-resident NK cells derive from precursors in the conventional NK cell lineage. We further show that the genetic absence of uterine NK cells during murine gestation leads to adverse pregnancy outcomes marked by reduced litter sizes and increased resorption rates. Collectively, our data underscore the pivotal role of uNK cells in pregnancy and offer novel insights into their lineage specification, revealing Eomesodermin as a crucial factor in their establishment.

## Introduction

Natural killer (NK) cells are key effectors of the innate immune system that play a myriad of roles in immunity, ranging from fighting infection to controlling malignant transformation^1^. Conventional NK (cNK) cells originate in the bone marrow and continuously circulate through the blood, from where they are recruited to sites of inflammation or infection^2,3^. While cNK cells are predominantly present in peripheral blood, NK cell subsets can also be found residing within tissues and do not circulate. Tissue-resident NK (trNK) cells seed non-lymphoid tissue early in life independent of inflammatory stimuli and are retained there, forming long-lived, tissue-specific lymphocyte populations with unique phenotypic and functional attributes^4^.

In naïve mice, we previously described trNK cells in the liver, skin, kidney and virgin uterus^5–7^. The tissue-resident nature of these trNK cells, as opposed to the circulating characteristic of cNK cells, was established by parabiosis^5,6,8^. trNK cells resemble cNK cells in terms of surface markers, such as NK1.1 and NKp46, and functions, including cytokine production and degranulation capacity^5–7^. However, trNK cells can be distinguished from cNK cells by the mutually exclusive expression of CD49a and CD49b. Specifically, trNK cells are CD49a^+^ CD49b^−^, while cNK cells are CD49a^−^ CD49b^+^^5–7^. Furthermore, trNK cells appear to follow distinct developmental trajectories compared to cNK cells as supported by their unique transcriptional requirements. For example, trNK cells in the liver and skin require the T-box transcription factor, T-bet, which has only modest effects on cNK cell development^6,9^. This dependence on T-bet makes trNK cells in the liver and skin phenotypically similar to classical type 1 innate lymphoid cells (ILC1s) rather than cNK cells^10,11^. Conversely, while NF-IL3 is crucial for the development of cNK cells, it plays no role in the development of trNK cells, emphasizing the distinct developmental pathways and unique lineage identities of these NK cell subsets^6,7,12–15^. Interestingly, neither T-bet nor NF-IL3 are required for the development of uterine trNK cells, highlighting the diverse regulatory mechanisms that govern the differentiation and function of trNK cell populations across different organs^6,16^. This transcriptional distinction sets uterine trNK cells apart from trNK cells in the liver and skin.

While NK cells share phenotypic markers with other innate lymphoid cells (ILCs), particularly ILC1s, there are key functional and developmental distinctions between these cell types^10,17^. ILC1s are tissue-resident innate lymphocytes characterized by the potent production of IFNγ and TNFα that play a frontline role in tissue homeostasis and host defense^10,17–20^. Like NK cells, ILC1s express the surface markers NK1.1 and NKp46 and largely produce IFNγ^10^. When other ILCs were identified, NK cells were initially classified as ILC1s due to the production of IFNγ. This suggested that NK cells and ILC1s might represent related subsets arising from a single lineage. However, recent classification schemes have separated NK cells from ILC1s given that NK cell cytotoxic function makes NK cells more analogous to cytotoxic CD8^+^ T cells while ILC1 cytokine secretion makes ILC1s more akin to CD4^+^ T helper cells^21^. Furthermore, detailed lineage tracing studies have indicated that NK cells differentiate from the early ILC progenitor (EILP) cell rather than the downstream common help-like ILC progenitor (CHILP), which is required for all ILC subset differentiation including ILC1s^11,22^. Lastly, the separation of NK cells from ILC1s is further supported by NK cell dependence on the T-box transcription factor, Eomesodermin (Eomes), which is dispensable for ILC1 development^9–11,23^. Together, thorough lineage studies and NK cell dependence on Eomes reinforce the notion that NK cells and ILC1s arise from distinct lineages, with NK cells representing a separate immune cell lineage from all ILCs.

Recent studies, however, have further blurred the relationship between ILC1s, cNK cells, and trNK cells. Under certain pathological conditions, cNK cells and ILC1s display developmental plasticity subsequent to their lineage specification, wherein cNK cells can convert into ILC1-like cells. In the tumor microenvironment, TGF-β signaling induces the conversion of cNK cells into ILC1-like cells to promote tumor immunoevasion^24^. Similarly, *in vitro* exposure to TGF-β drives splenic cNK cells to adopt a ILC1-like phenotype resembling that of salivary gland ILC1s ^25^. In the obese murine liver, cNK cells shift toward a less cytotoxic, ILC1-like phenotype to protect against non-alcoholic fatty liver disease^26^. Lastly, *Toxoplasma gondii* infection drives the conversion of cNK cells into ILC1-like cells that are phenotypically distinct from both cNK cells and ILC1s at homeostatic conditions^27^. Since trNK cells greatly resemble ILC1s, it has become difficult to determine if trNK cells form a population of cells distinct from ILC1s or represent interrelated subsets that adopt unique phenotypes in certain microenvironments.

During pregnancy, uterine NK (uNK) cells are the most abundant leukocytes at the maternal-fetal interface in both mice and humans^28–31^. The NK cell population in the pregnant uterus is heterogenous, consisting primarily of trNK cells and few cNK cells^6,28,32^. Unlike cNK cells in peripheral circulation, uNK cells are poorly cytotoxic but still capable of producing cytokines^33–35^. Due to their unique phenotype in the pregnant uterus, uNK cells have been thought to contribute to various physiological aspects of gestation that extend beyond host-defense, including placental angiogenesis, fetal trophoblast differentiation, and the production of growth factors supporting fetal development, though it has been difficult to demonstrate that uNK cells, in the absence of defects in other immune cells, affect fecundity^35–42^. Despite their apparent critical roles in pregnancy, the mechanisms by which uNK cells arise as well as the precise contributions of uNK cells to successful pregnancy outcomes remain elusive.

In this study, we found that Eomes is a key transcription factor driving the establishment of trNK cells in both the virgin and pregnant uterus. Furthermore, we demonstrate that the selective loss of uNK cells during murine gestation results in suboptimal reproductive outcomes characterized by markedly reduced litter sizes and increased resorption rates.

## Materials and Methods

### Mice

All mouse studies were conducted in accordance with ethical guidelines and animal protocol approved by the Washington University School of Medicine Animal Studies Committee under protocol number 21-0090. Wild-type *C57BL/6* (stock number 665) mice and *B6.SJL-Ptprc^a^Pepc^b^/BoyJ* (stock number 664) mice were purchased from Charles River Laboratories (Wilmington, MA). *Ncr1^iCre^* mice were kindly gifted by Eric Vivier at Aix Marseille University, Marseille, France. Eomes^Ncr^^1^^Δ^ mice were generated by crossing *B6.129S1(Cg)-Eomes^tm^*^1^*^.1Bflu^/J* (strain: 017293; The Jackson Laboratory, Bar Harbor, ME) with *Ncr1*^i*Cre*^ mice. *Rag2*^-/-^γ*C*^-/-^ mice were generated by crossing *B6.Cg-Rag2^tm^*^1^*^.1Cgn^/J* mice (strain 008449; The Jackson Laboratory) with *B6.129S4-Il2rg^tm1Wjl^/J* mice (strain 003174; The Jackson Laboratory) as previously described^27^. Female mice from 7-10 weeks of age were used in all mouse studies. Soiled cages were swapped amongst mice of different genotypes for at least two weeks to expose all mice in the same experiment to the same microbiome. All mice were housed in the Laboratory for Animal Care barrier facility at the Washington University School of Medicine.

### Timed-Pregnancies

Virgin C57BL/6, littermate control, Rag2^-/-^γC^-/-^, and Eomes^Ncr^^1^^Δ^ virgin female mice were mated with C57BL/6-CD45.1 male mice overnight. Female mice were mated with C57BL/6 CD45.1 male mice to distinguish maternal lymphocytes from fetal lymphocytes at the maternal-fetal interface. Timing of conception was determined by detection of copulation plug the following morning. Presence of copulation plug was designated as gestational day (gd) 0.5. Pregnancy outcomes were evaluated by comparing litter sizes, pup weight at birth, and gestational length at first parturition between C57BL/6, littermate, Rag2^-/-^γC^-/-^, and Eomes^Ncr^^1^^Δ^ dams. Pregnant dams were dissected at either gds 6.5 and 9.5 to assess the immune constituents of implantation sites, or at gds 9.5 and 13.5 to assess resorption rates. Resorption rates were calculated as the percentage of resorbed implantation sites per pregnant uterus.

### Single Cell Suspension Preparation from Different Tissues

#### Implantation Site Digestion

All healthy implantation sites were harvested from pregnant dams at gds 6.5 and 9.5 and prepared for flow cytometric analysis. Each pregnant uterus was digested with Liberase TL (167 µg/ml; Sigma-Aldrich, St. Louis, MO) and DNase1 (150 μg/ml; Sigma-Aldrich) for 1 hour at 37°C. Digested implantation sites were minced, washed with 10% FBS RPMI media, and resuspended in 2mL of complete R10 media. We ensured that all surface markers of interest remained unaffected by the digestion enzymes used in these studies by spiking the preparations with congenically marked splenocytes for comparison to untreated splenocytes by flow cytometry.

#### Liver Preparation

Livers from each pregnant dam were harvested, minced, and filtered through a 100μM mesh. Liver suspensions were subsequently treated with Percoll and 10% 10X PBS to isolate hepatic lymphocytes, which were then treated with RBC Lysis, washed with R10 media, and resuspended in 200 μL of complete R10 media.

#### Splenic Preparation

Spleens from every pregnant dam were harvested, minced, and filtered through a 70μΜ mesh. Splenocyte suspensions were subsequently treated with RBC Lysis, washed with 10% FBS RPMI media, and resuspended in 5 mL of complete R10 media.

### Flow Cytometry

Fluorescent-labeled antibodies purchased from Invitrogen (Waltham, MA) including CD3e (clone 145-2C11), CD19 (1D3), CD45.2 (104), CD45.1 (A20), CD49b (DX5), EOMES (Dan11mag), Ly49H (3D10), Ly49I (YLI-90), NKG2AB6 (16a11), NKp46 (29A1.4), and Fixable Viability Dye (eFluor^TM^506); BioLegend (San Diego, CA) including CD200R (OX-110), CXCR6 (SA051D1), Ly49A (YE1/48.10.6), and NK1.1 (PK136); eBioscience (San Diego, CA) including CD69 (H1.2F3), Ki67 (SolA15), and KLRG1 (2F1); and BD Biosciences (Franklin Lakes, NJ) including CD49a (Ha31/8).

For quantification of cell numbers, 5,000 Precision Count Beads (BioLegend) were added prior to staining. Cells were first stained with fixable viability dye (Invitrogen) and subsequently stained for cell surface molecules in 2.4G2 hybridoma supernatant (anti-FcγRIII) to ensure that Fc receptors were blocked. Afterwards, cells were fixed and permeabilized using the Foxp3/Transcription Factor Staining Buffer Set (eBiosciences) and then stained for intracellular molecules. All samples were acquired on a FACS Canto (BD Bioscience) and analyzed using FlowJO software (BD Biosciences). Maternal CD49b^+^ Eomes^+^ cNK cells were defined as viable singlets, CD3^-^ CD19^-^ CD45.1^-^ CD45.2^+^ NK.1^+^ NKp46^+^ CD49a^-^ CD49b^+^ Eomes^+^. Maternal CD49a^+^ Eomes^+^ trNK cells were defined as viable singlets, CD3^-^ CD19^-^ CD45.1^-^ CD45.2^+^ NK.1^+^ NKp46^+^ CD49a^+^ CD49b^-^ Eomes^+^. Maternal CD49a^+^ Eomes^-^ ILC1s were defined as viable singlets CD3^-^ CD19^-^ CD45.1^-^ CD45.2^+^ NK.1^+^ NKp46^+^ CD49a^+^ CD49b^-^ Eomes^-^.

### Statistical Analysis

Statistical analysis was performed with Prism (GraphPad software) using unpaired *t* tests and one-way ANOVA corrected for multiple comparisons as indicated in the figure legends. Normality was with Prism (GraphPad software) using Q-Q plots. Error bars in figures represent the SEM. Statistical significance was indicated as follows: ns, not significant; \**p* < 0.05; ** *p* < 0.01; *** *p* < 0.001; and **** *p* < 0.0001.

## Results

### CD49a^+^ Eomes^+^ NK cells in the virgin and pregnant uterus are Eomes-dependent

Consistent with prior studies, analysis of the ILC subsets in the virgin uterus from C57BL/6 female mice revealed three distinct ILC subsets: CD49a^+^ Eomes^+^ trNK cells, CD49a^+^ Eomes^-^ ILC1s, and CD49b^+^ Eomes^+^ cNK cells (**Figure 1 C and D**)^32^. We then aimed to confirm the presence of these three ILC subsets within the pregnant uterus. In our studies, virgin female mice were mated with C57BL/6 CD45.1 male mice to distinguish maternal lymphocytes from fetal lymphocytes at the maternal-fetal interface^38,43–45^. As expected, the three distinct ILC subsets were also observed in the pregnant uterus during early gestation (**Figure 1 E-G**). The phenotypic characterization of these ILC subsets was consistent with prior studies, supporting the prevailing notion that uterine CD49a^+^ Eomes^+^ trNK cells may represent a unique subset of tissue-resident innate immune cells distinct from ILC1s (**Figure 2**)^8,32,46^.

**Figure 1.**
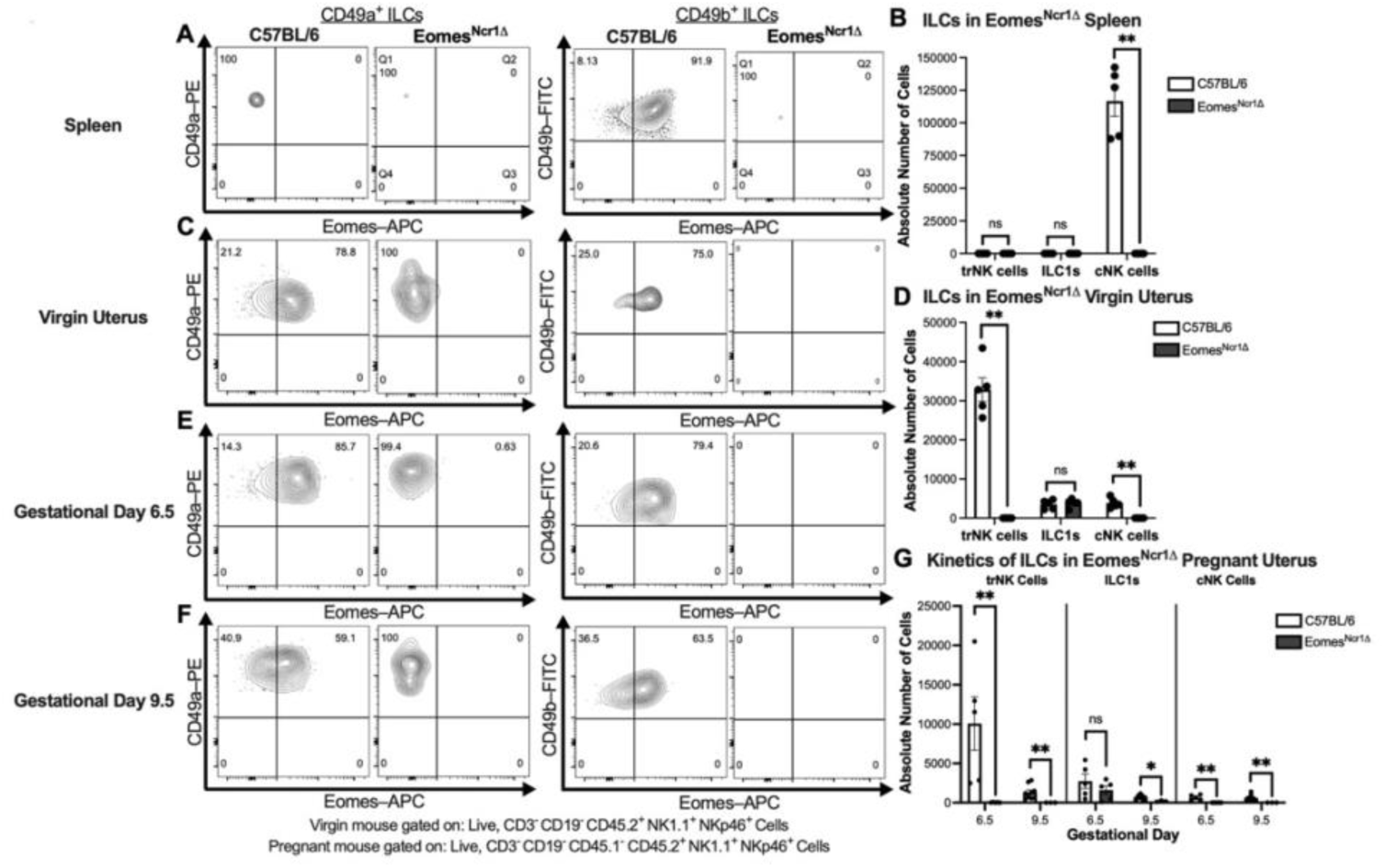
Loss of *Eomesodermin* expression in *Ncr1*-expressing cells ablates CD49a^+^ Eomes^+^ NK cells in the virgin and pregnant uterus. (**A**) Representative flow plots showing the expression of CD49a, CD49b, and Eomes across ILC subsets in the spleen of C57BL/6 and Eomes^Ncr1Δ^ female mice (C57BL/6, *n*=5, Eomes^Ncr1Δ^, *n*=5). (**B**) Absolute cell counts of CD49a^+^ Eomes^+^ trNK cells, CD49a^+^ Eomes^-^ ILC1s, and CD49b^+^ Eomes^+^ cNK cells in the spleen of C57BL/6 and Eomes^Ncr1Δ^ female mice. (C) Representative flow plots showing the expression of CD49a, CD49b, and Eomes across ILC subsets in the virgin uterus of C57BL/6 and Eomes^Ncr1Δ^ female mice (C57BL/6, *n*=5, Eomes^Ncr1Δ^, *n*=5). (D) Absolute cell counts of CD49a^+^ Eomes^+^ trNK cells, CD49a^+^ Eomes^-^ ILC1s, and CD49b^+^ Eomes^+^ cNK cells in the virgin uterus of C57BL/6 and Eomes^Ncr1Δ^ female mice. (**E**) Representative flow plots showing the expression of CD49a, CD49b, and Eomes across ILC subsets in the pregnant uterus of C57BL/6 and Eomes^Ncr1Δ^ dams at gd 6.5 (C57BL/6, *n*=5, Eomes^Ncr1Δ^, *n*=5). (**F**) Representative flow plots showing the expression of CD49a, CD49b, and Eomes across ILC subsets in the pregnant uterus of C57BL/6 and Eomes^Ncr1Δ^ dams at gd 9.5 (C57BL/6, *n*=9, Eomes^Ncr1Δ^, *n*=3). (**G**) Absolute cell counts of CD49a^+^ Eomes^+^ NK cells, CD49a^+^ Eomes^-^ cell, and CD49b^+^ Eomes^+^ cells in the pregnant uterus of C57BL/6 and Eomes^Ncr1Δ^ dams at gd 6.5 and gd 9.5 (gd 6.5: C57BL/6, *n*=5, Eomes^Ncr1Δ^, *n*=5; gd 9.5: C57BL/6, *n*=9, Eomes^Ncr1Δ^*, n*=3). Virgin uterine samples gated on Live, CD3^-^ CD19^-^ CD45.2^+^ NK1.1^+^ NKp46^+^ cells. Pregnant uterine samples gated on Live, CD3^-^ CD19^-^ CD45.1^-^ CD45.2^+^ NK1.1^+^ NKp46^+^ cells. Statistics were calculated unpaired *t tests* with the Mann-Whitney correction. Error bars indicate SEM; ns, not significant; ^✱^*p* < 0.05; ^✱✱^*p* <0.01.

**Figure 2.**
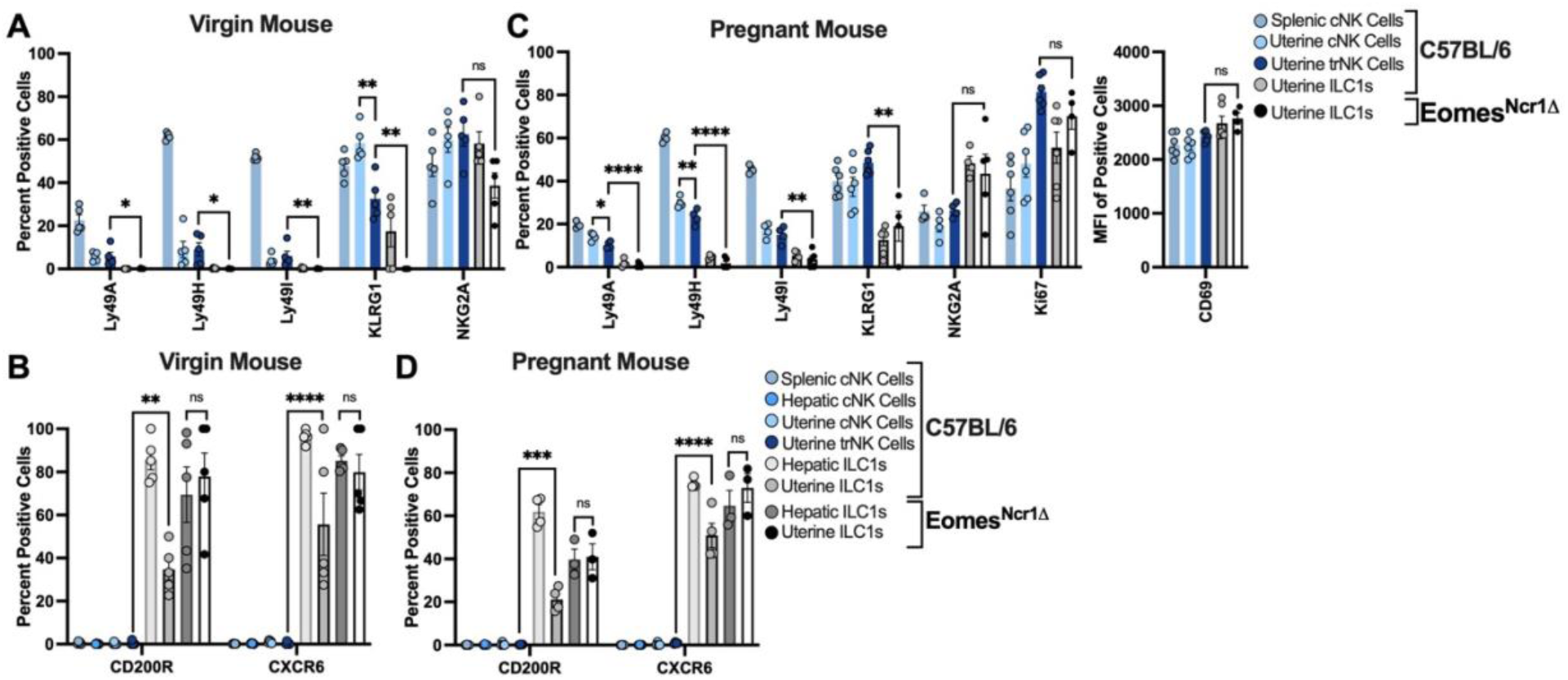
Phenotypic characterization of ILC subsets in the virgin and pregnant uterus of Eomes^Ncr1Δ^ dams. (**A**) Receptor repertoire of splenic and uterine cNK, trNK, and ILC1s subsets in C57BL/6 and Eomes^Ncr^^1^ ^Δ^ virgin female mice (C57BL/6, *n*=5, Eomes^Ncr1Δ^, *n*=5). (**B**) Expression of CD200R and CXCR6 of splenic, hepatic, and uterine cNK, trNK, and ILC1s subsets in C57BL/6 and Eomes^Ncr1Δ^ virgin female mice (C57BL/6, *n*=5, Eomes^Ncr1Δ^, *n*=5. (**C**) Receptor repertoire of splenic and uterine cNK, trNK, and ILC1s subsets in C57BL/6 and Eomes^Ncr1Δ^ pregnant dams at gd 6.5 (C57BL/6, *n*=5, Eomes^Ncr1Δ^, *n*=4–5). (**D**) Expression of CD200R and CXCR6 of splenic, hepatic, and uterine cNK, trNK, and ILC1s subsets in C57BL/6 and Eomes^Ncr1Δ^ pregnant dams at gd 6.5 (C57BL/6, *n*=4, Eomes^Ncr1Δ^, *n*=3). Virgin uterine samples gated on Live, CD3^-^ CD19^-^ CD45.2^+^ NK1.1^+^ NKp46^+^ cells. Pregnant uterine samples gated on Live, CD3^-^ CD19^-^ CD45.1^-^ CD45.2^+^ NK1.1^+^ NKp46^+^ cells. Statistics were calculated with an Ordinary one-way ANOVA corrected for multiple comparisons. Normality was confirmed using Q-Q plots. Error bars indicate SEM; ns, not significant; ^✱^*p* < 0.05, ^✱✱^*p* <0.01, ^✱✱✱^*p* <0.001, ^✱✱✱✱^ *p* <0.0001. MFI, median fluorescent intensity.

Next, we sought to determine whether Eomes affects the establishment of trNK cells in the virgin and pregnant uterus. Briefly, *Eomes^fl/fl^* mice were crossed with *Ncr1^icre^* mice to generate Eomes^Ncr^^1^^Δ^ mice. In accordance with the literature, cNK cells were markedly reduced from the spleen, virgin uterus, and pregnant uterus of Eomes^Ncr^^1^^Δ^ dams (**Figure 1A, C, E, F**)^9,23^. Surprisingly, the loss of Eomes also completely ablated the population of CD49a^+^ Eomes^+^ trNK cells in the virgin uterus as well as in implantation sites in Eomes^Ncr^^1^^Δ^ female mice (**Figure 1A, C, E, F)**. The numbers of CD49a^+^ Eomes^-^ ILC1s in the virgin and pregnant uterus were unaffected by the loss of Eomes expression in *Ncr1*-expressing cells (**Figure 1D, G**). Taken together, these results establish that trNK cells in the uterus are dependent on Eomes, suggesting these cells differentiate from progenitors in the cNK cell lineage.

### Uterine CD49a^+^ Eomes^-^ ILC1s in Eomes^Ncr^^1^^Δ^ female mice resemble bona fide ILC1s at steady-state and during early gestation

Eomes is essential for the maturation of cNK cells, as the loss of Eomes expression from mature cNK cells has been shown to induce a reversion into a phenotypically immature state^23^. This raises the possibility that the residual population of CD49a^+^ Eomes^-^ cells in the virgin and pregnant uterus of Eomes^Ncr^^1^^Δ^ dams may represent an immature NK cell population. Therefore, we questioned whether this residual population exhibited a phenotypic profile consistent with classical ILC1s.

Like traditional ILC1s in the virgin uterus of C57BL/6 female mice, CD49a^+^ Eomes^-^ ILC1s in the virgin uterus of Eomes^Ncr^^1^^Δ^ female mice did not express Ly49A, Ly49H, Ly49I, and KLRG1 but showed moderate expression of NKG2A (**Figure 2A and Supplemental Figure 1**). Notably, CD49a^+^ Eomes^-^ ILC1s in the virgin uterus of Eomes^Ncr^^1^^Δ^ female mice exhibited robust expression of both CD200R and CXCR6, makers classically associated with ILC1s (**Figure 2B**). Similarly, the residual population of CD49a^+^ Eomes^-^ ILC1s in the pregnant uterus of Eomes^Ncr^^1^^Δ^ dams lacked expression of traditional cNK cells markers but displayed high expression of CD200R and CXCR6 (**Figure 2C** and **D**). Additionally, uterine CD49a^+^ Eomes^-^ ILC1s in Eomes^Ncr^^1^^Δ^ pregnant dams expressed Ki67 and CD69, suggesting these cells were activated by the physiological state of pregnancy (**Figure 2C**). Collectively, these findings suggest that residual CD49a^+^ Eomes^-^ ILC1s in the virgin and pregnant uterus of Eomes^Ncr^^1^^Δ^ female mice are bona fide ILC1s, indicating that loss of Eomes in *Ncr1*-expressing cells does not affect the population of uterine ILC1s in Eomes^Ncr^^1^^Δ^ female mice.

### Loss of uNK cells results in adverse pregnancy outcomes

The selective loss of uNK cells upon the deletion of Eomes expression in all Ncr1-expressing cells provided the unique opportunity to evaluate the precise contributions of uNK cells to successful pregnancy outcomes. Gestational length, litter size, and pup birth weight were compared between C57BL/6, littermate control, Rag2^-/-^γC^-/-^, and Eomes^Ncr^^1^^Δ^ dams mated with C57BL/6 CD45.1 males. The loss of NK cells in the pregnant uterus markedly reduced litter sizes in Eomes^Ncr^^1^^Δ^ dams compared to both C57BL/6 and littermate control dams (**Figure 3A**). This finding was further supported by an increased resorption rate in Eomes^Ncr^^1^^Δ^ dams at gd 9.5 (**Figure 3D)**. Moreover, the increased resorption rate observed in Eomes^Ncr^^1^^Δ^ dams at the start of mid-gestation correlates with the rise and decline of uterine CD49a^+^ Eomes^+^ NK cell numbers by mid-gestation. The pregnancy abnormalities observed in Eomes^Ncr^^1^^Δ^ dams mirrored those in Rag2^-/-^γC^-/-^ dams, reinforcing an indispensable role for uNK cells in pregnancy. However, birth weight of neonatal pups was not affected by the loss of uNK cells, and no correlations were discerned between litter size and pup birth weight (**Figure 3B**). Furthermore, there was no notable difference in the mean gestational length in the absence of uNK cells as the gestational period of Eomes^Ncr^^1^^Δ^ dams was statistically indistinguishable from that of C57BL/6, littermate control, and Rag2^-/-^γC^-/-^ dams (**Figure 3C**). The lack of differences in pup birth weight and gestational period suggests that the increase in fetal demise observed in Eomes^Ncr^^1^^Δ^ dams is as a stochastic, all-or-nothing event. This implies that in the absence of uNK cells, pregnancies either proceed to term with normal fetal development or experience complete loss. Regardless, taken together our results demonstrate that pregnancy outcomes are significantly compromised in the absence of uNK cells.

**Figure 3.**
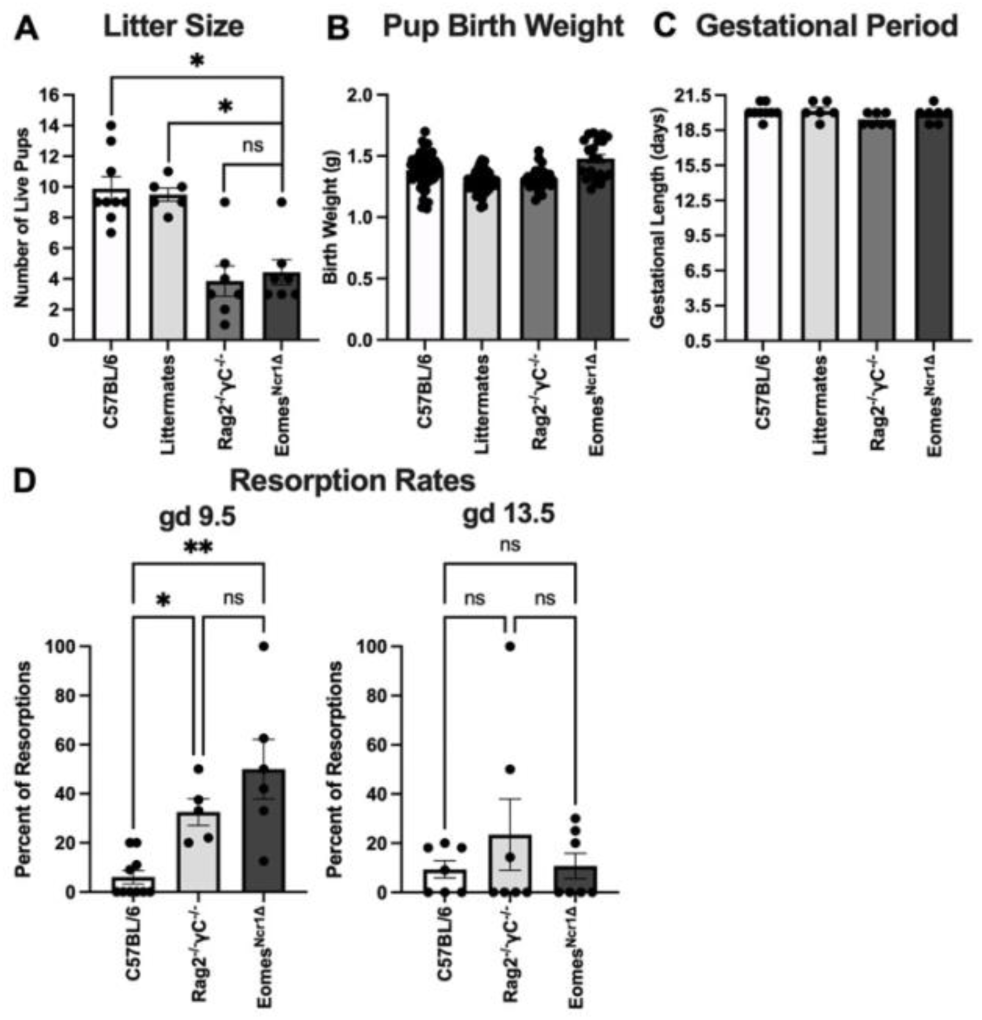
Loss of uNK cells in the pregnant uterus reduced litter sizes and increased resorptions rates. **(A)** Number of live pups at first parturition from C57BL/6, littermate control, Rag2^-/-^γC^-/-^, and Eomes^Ncr1Δ^ dams (C57BL/6, *n*=9, Littermates, *n*=6, Rag2^-/-^γC^-/-^, *n*=7, Eomes^Ncr1Δ^, *n*=7). (**B**) Pup birth weight in grams (g) from pups birthed by C57BL/6, littermate control, Rag2^-/-^γC^-/-^, and Eomes^Ncr1Δ^ dams (C57BL/6, *n*=78, Littermates, *n*=67, Rag2^-/-^γC^-/-^, *n*=27, Eomes^Ncr1Δ^, *n*=22). (**C**) Gestational period in days for C57BL/6, littermate control, Rag2^-/-^γC^-/-^, and Eomes^Ncr1Δ^ dams (C57BL/6, *n*=9, Littermates, *n*=6, Rag2^-/-^γC^-/-^, *n*=7, Eomes^Ncr1Δ^, *n*=7). **(D)** Eomes^Ncr1Δ^ dams exhibited a higher percentage of resorbed implantation sites at gd 9.5 (gd 9.5: C57BL/6, *n*=10, Rag2^-/-^γC^-/-^, *n*=5, Eomes^Ncr^^1^ ^Δ^, *n*=6; gd 13.5: C57BL/6, *n*=7, Rag2^-/-^γC^-/-^, *n*=7, Eomes^Ncr1Δ^, *n*=7). Resorption rates (RR) were calculated as: RR(%) = (number of resorbed implantation sites/number of total implantation sites) X 100. Statistics were calculated using Kruskal-Wallis test corrected for multiple comparisons. Error bars indicate SEM; ns, not significant; ^✱^*p*< 0.05 and ^✱✱^*p* <0.01.

## Discussion

Here, we demonstrate that Eomes is a lineage-determining transcription factor for trNK cells in the uterine microenvironment. Although Eomes is known to regulate the development and maturation of cNK cells, a role for Eomes as a key developmental driver for uterine trNK cells had not been previously described. Moreover, to our knowledge, our study is the first to demonstrate that the loss of uNK cells by genetic means in the pregnant uterus results in adverse pregnancy outcomes characterized by markedly reduced litter sizes and increased resorption rates. Taken together, these data not only shed light on a key transcriptional factor underpinning the establishment of uterine trNK cells but also provide direct genetic evidence supporting a central role for uNK cells during gestation.

Recent studies have significantly broadened our understanding of trNK cells across various organs. However, uterine trNK cells remain largely understudied despite evidence linking disruptions in uNK cells with adverse pregnancy outcomes in immunodeficient mouse models^16,34,36,47,48^. Furthermore, there was no previous consensus on the developmental origins of uterine trNK cells. Prior studies have shown the number of uterine trNK cells was unaffected in in T-bet-deficient mice, suggesting that trNK cells may not differentiate from the ILC1 lineage^6^. Moreover, peripheral cNK cells can be distinguished from ILC1s by their dependance on IL-15rα for survival^49–53^, while all uNK cell subsets are ablated in both IL-15 and IL-15rα-deficient mice, suggesting that uterine trNK cells are more related to cNK cells than ILC1s^6,54,55^. Indeed, given that Eomes is a distinguishing transcription factor that differentiates cNK cells from ILC1s, our data supports the notion that uterine trNK cells differentiate from precursors of the cNK cell lineage. New literature demonstrates that peripheral cNK cells can differentiate into uterine trNK cells in a progesterone-dependent manner, further supporting that the loss of uterine trNK cells could be attributed to the loss of cNK cells in the periphery upon the deletion of Eomes expression in all Ncr1-expressing cells^56^. However, unlike cNK cells which require NF-IL3 for their development, uterine trNK cells have been found to develop independently of NF-IL3^6,13–16,57^. The presence of uterine trNK cells in NF-IL3-deficient mice suggested that uterine trNK cells may differentiate from a distinct lineage than cNK cells since NF-IL3 expression is induced during the early stages of cNK cell development^13,15^. The persistence of uterine trNK cells in the absence of NF-IL3 expression may be explained by the possible involvement of yet to be defined cNK cell precursors or other transcription factors that influence the early developmental stages of cNK cells. Previously, the developmental stage at which Eomes enforces commitment to the NK cell lineage remained unclear; however, a recent study demonstrates Eomes expression defines the early bone marrow progenitor to cNK cells in the mouse^58^. Therefore, Eomes may be required at an early stage of cNK cell development, giving rise to a precursor that yields cNK cells as well as uterine trNK cells. NF-IL3 may then function downstream and affect a precursor that is required for further cNK cells differentiation but is not required for uterine trNK cells.

The dependence of uterine trNK cells on Eomes resembles similar processes in other tissue-specific ILC populations. In the salivary glands, TGF-β suppresses Eomes, promoting the differentiation of ILC1-like cells^25^. A similar transformation occurs in the tumor microenvironment, where upon exposure to TGF-β, cNK cells downregulate Eomes and upregulate T-bet, adopting ILC1-like features^24^. During *Toxoplasma gondii* infection, this process is recapitulated, with cNK cells downregulating Eomes and differentiating into ILC1-like cells. Notably, this phenotypic shift is dependent on T-bet expression, as it is lost in T-bet-deficient mice. Interestingly, a subset of Eomes^+^ CD49a^+^ ILC1-like cells, which resemble uterine trNK cells, arises in both wild-type and T-bet-deficient mice during *Toxoplasma gondii* infection^27^. This plasticity, driven by the interplay between Eomes and T-bet, highlights how environmental cues shape immune cell function across different tissues. For uterine trNK cells, however, this plasticity appears to be a normal physiological response, more akin to salivary gland differentiation of NK cells into ILC1-like cells.

Emerging studies suggest that Eomes plays distinct roles in the formation and maintenance of tissue-resident immune cells across different tissues. For example, in the liver, kidney, and skin, Eomes represses the formation of tissue-resident memory T-cells^59,60^. However, in the intestine, while Eomes is dispensable for the formation of tissue-resident memory T-cells, it is essential for their maintenance^61^. The context-dependent role of Eomes in the formation and maintenance of tissue-resident immune cell populations suggests the factors required for the formation of these cells may differ from those required for their maintenance. While our study ascertains a role for Eomes in the establishment of uterine trNK cells, additional studies are needed to identify the factors regulating their maintenance in the uterine microenvironment.

The exact contributions of uNK cells to pregnancy success remain incompletely understood. Current insights into the roles of uterine trNK cells during pregnancy come from studies investigating pregnancy outcomes across different immunodeficient mouse models. Pioneering studies demonstrated that NK cell reconstitution reversed the reproductive and placentation defects observed in Tgε26 pregnant dams^47,48^. The involvement of uNK cells in decidual angiogenesis has been inferred from histological comparisons of implantation sites from BALB/c^+/+^, Rag2^-/-^, Rag2^-/-^γC^-/-^, and Tgε26 dams^34,36^. Additionally, the development of the NF-IL3-deficient mouse linked perturbations in uNK cell subsets with poor decidualization and improper placental vascularization during early gestation^16^. In addition, uNK cells promote fetal growth by secreting growth factors^37^. In humans, decidual CD49a^+^ Eomes^+^ NK cells promote fetal development during early pregnancy, and their reduced numbers are associated with recurrent spontaneous abortion^40,62,63^. Together, these studies imply that disruptions in the uNK cell population are associated with serious reproductive phenotypes and adverse pregnancy outcomes. However, since the contributions of uNK cells to pregnancy have been deduced using mouse models that lack isolated NK cell deficiencies, the results of these studies did not constitute direct evidence of the specific ways uNK cells promote successful reproductive outcomes. Our study fills this gap by using a mouse model with an isolated uNK cell deficiency showing adverse pregnancy outcomes. Additional studies are necessary to pinpoint the precise roles of uNK cells in gestation and will be aided by further studies of these mice.

## Disclosures

The authors have no financial conflicts of interest.

## Acknowledgements

We extend our gratitude to the members of the Yokoyama Laboratory for insightful and valuable discussions.

## Footnotes

**Abbreviations used in this paper include**
cNK: conventional NK
Eomes: Eomesodermin
gd: gestational day
ILC: innate lymphoid cell
ILC1: type 1innate lymphoid cell
NK: natural killer
trNK: tissue-resident NK
uNK: uterine NK

## Notes

### Competing Interest Statement

The authors have declared no competing interest.

