## Supplemental Figure 1 for "Eomesodermin defines uterine NK cells crucial for pregnancy success in mice"

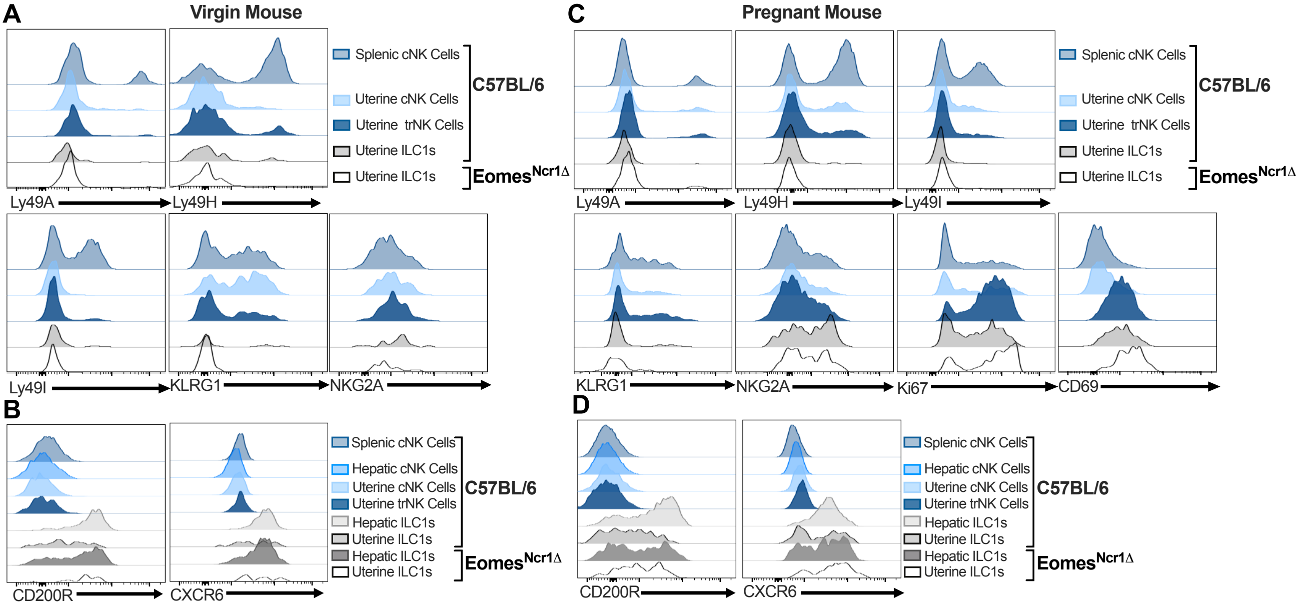


**Supplemental Figure 1. Receptor Repertoire of ILC subsets in the virgin and pregnant uterus of Eomes$^{Ncr1\Delta}$ dams.** (**A**) Concatenated histograms depicting the receptor repertoire of splenic and uterine cNK, trNK, and ILC1s subsets in C57BL/6 and Eomes$^{Ncr1\ \Delta}$ virgin female mice (C57BL/6, *n*=5, Eomes$^{Ncr1\Delta}$, *n*=5). (**B**) Concatenated histograms depicting the expression of CD200R and CXCR6 of splenic, hepatic, and uterine cNK, trNK, and ILC1s subsets in C57BL/6 and Eomes$^{Ncr1\Delta}$ virgin female mice (C57BL/6, *n*=5, Eomes$^{Ncr1\Delta}$, *n*=5. (**C**) Concatenated histograms depicting the receptor repertoire of splenic and uterine cNK, trNK, and ILC1s subsets in C57BL/6 and Eomes$^{Ncr1\Delta}$ pregnant dams at gd 6.5 (C57BL/6, *n*=5, Eomes$^{Ncr1\Delta}$, *n*=4–5). (**D**) Concatenated histograms depicting the expression of CD200R and CXCR6 of splenic, hepatic, and uterine cNK, trNK, and ILC1s subsets in C57BL/6 and Eomes$^{Ncr1\Delta}$ pregnant dams at gd 6.5 (C57BL/6, *n*=4, Eomes$^{Ncr1\Delta}$, *n*=3). Virgin uterine samples gated on Live, CD3$^-$ CD19$^-$ CD45.2$^+$ NK1.1$^+$ NKp46$^+$ cells. Pregnant uterine samples gated on Live, CD3$^-$ CD19$^-$ CD45.1$^-$ CD45.2$^+$ NK1.1$^+$ NKp46$^+$ cells.
